# Loss of HMGCS2-mediated intestinal stem cell ketogenesis is a metabolic barrier to mucosal healing in Ulcerative colitis

**DOI:** 10.1101/2025.08.25.671702

**Authors:** Duncan G Rutherford, Verity E Cowell, Robert J Whelan, Alexandra J B Cavanagh, Claire E Adams, Peter D Cartlidge, Phoebe Y Lau, Mairi H I Hodge, Nigel B Jamieson, Claire K Dietrich, Darren Ruane, Joshua Rusbuldt, Koichi Hashikawa, Anna Smith, Laura C Greaves, Gareth-Rhys Jones, Gwo-tzer Ho

## Abstract

Human colonic epithelial cells express high levels of 3-Hydroxy-3-methylglutaryl-CoA synthase 2 (HMGCS2), the crucial mitochondrial enzyme responsible for the production ketone bodies, predominantly beta-hydroxybutyrate (βHB). Ketogenesis is an important metabolic pathway responsible for energy production in states of fasting and in the intestine ketone bodies are also important in regulating intestinal stem cell (ISC) homeostasis and regeneration. Using combined single-cell and spatial transcriptomics, organoids and prospective longitudinal mucosal analysis, we demonstrated a profound loss of ISC HMGCS2-mediated ketogenesis with consequential detrimental effect on Ulcerative colitis (UC), a chronic inflammatory bowel disease that affects ∼4 million individuals worldwide. We show that βHB restores UC ISC metabolic function, reduces cellular stress with further evidence of epigenetic programming key for restoration of ISC function. Furthermore, low colonic *HMGCS2* expression is associated with treatment failure of multiple biological immune therapies in UC. Our findings reveal the crucial role for HMGCS2 and loss of ketogenesis as the ‘tipping point’ for pathogenic epithelial dysfunction and importantly, a promising metabolic therapeutic target for UC.

## Introduction

Ulcerative colitis (UC) is a chronic inflammatory bowel disease (IBD) with characteristic confluent, superficial, immune-mediated inflammation of the colon^1^. Beyond an aberrant immunologic host response, colonic epithelial dysfunction is a major component of UC pathogenesis^2,3^. In health, epithelial integrity is maintained by highly metabolically active intestinal stem cells (ISCs), situated at the base of colonic crypts that replenish the colonic epithelium every 3-5 days^4^. In UC, intestinal inflammation is associated with poor mucosal healing despite potent advanced immune therapies. Several lines of evidence have shown mitochondrial damage within colonic epithelial cells compromises its function^5–7^. Within the colon, mitochondria are particularly exposed to noxious stimuli, compounded by loss of protective mechanisms in disease states^8,9^. Metabolic alterations have broad reaching consequences for both cell fate determination and cellular function^10^, however little is known about ISC metabolism in UC and its impact on epithelial repair. Here, we identify defective ketogenesis as the metabolic ‘tipping point’ where mitochondria and metabolic abnormalities converge on ISC dysfunction in UC. Our data identifies mitochondrial-specific *3-Hydroxy-3-methylglutaryl-CoA synthase 2* (*HMGCS2*), the gene encoding the rate-limiting step for ketogenesis as the critical nexus. Importantly, this reveals a novel metabolic therapeutic avenue in UC.

### Intestinal stem cell mitochondrial and metabolic dysfunction in UC

To understand the behaviour of epithelial cells in a human disease and tissue specific context, we first employed a single-cell RNA sequencing (scRNAseq) of epithelial cells from endoscopically targeted colonic mucosal biopsies of treatment naïve newly diagnosed UC individuals and healthy controls (Fig. 1a-c, Extended Data. 1a and Supplementary Table 1). Within this dataset we identified 11 epithelial cell populations, with each patient sampled contributing to cell sub-types (Fig 1c). Firstly, we performed unsupervised differentially expressed gene (DEG) analysis of all *EPCAM*^+^ cells (UC inflamed vs control), which highlighted metabolic genes amongst the most significantly differentially regulated in UC (Fig. 1d) including *DUOXA2*, *ATP5PO*, *PPARGC1*α. To ascertain whether this was driven by specific epithelial cell populations, we next assessed metabolic pathway activity within each cell subtype. Here, we performed gene set enrichment analyses (GSEA) on DEGs between epithelial sub-types, observing that perturbations of metabolic pathways in ISCs and colonocytes were the most metabolically dysregulated (tricarboxylic acid (TCA) cycle, Oxidative phosphorylation [OXPHOS], Fatty acid metabolism) (Extended Fig. 1b). Specifically, within ISCs we observed downregulation of mitochondrial metabolic pathways (OXPHOS, TCA cycle) and upregulation of accessory metabolic pathways (alpha linolenic, arginine, glycolysis) (Fig. 1e). Furthermore, closer inspection of mitochondrial gene expression in UC ISCs (vs control) revealed upregulation of loci pertaining to mitochondrial stress (*VDAC3* and *HSPD1*), fission (*DNM1L* and *MFF*) and fusion (*MFN1* and *OPA1)* with downregulation of genes related to mitochondrial biogenesis and mitophagy (*PINK1* and *PPARGC1*α) (Fig. 1f). In line with transcriptomic changes, mitochondrial damage was evident by the loss of Complex I and IV proteins of the mitochondrial electron transport chain in colonic epithelial cells from UC patients with active disease (Fig. 1g, h Extended Fig. 1c, d and Supplementary Table 2). To gain further resolution of metabolic changes observed in the UC epithelium we generated a parallel colonic epithelial single-cell spatial transcriptomic (CosMx) dataset, finding downregulation of OXPHOS gene module and an associated compensatory upregulation of glycolysis (Fig. 1i, j, Extended Figure 1e and Supplementary Table 3). Collectively, these data reveal profound deregulation of mitochondrial and metabolic function within both the intestinal stem cells (ISCs) and the broader colonic epithelium in UC. Notably, the disruption within ISCs is particularly significant, as ISC function governs the downstream epithelial phenotype during differentiation and regeneration.

**Fig. 1.**
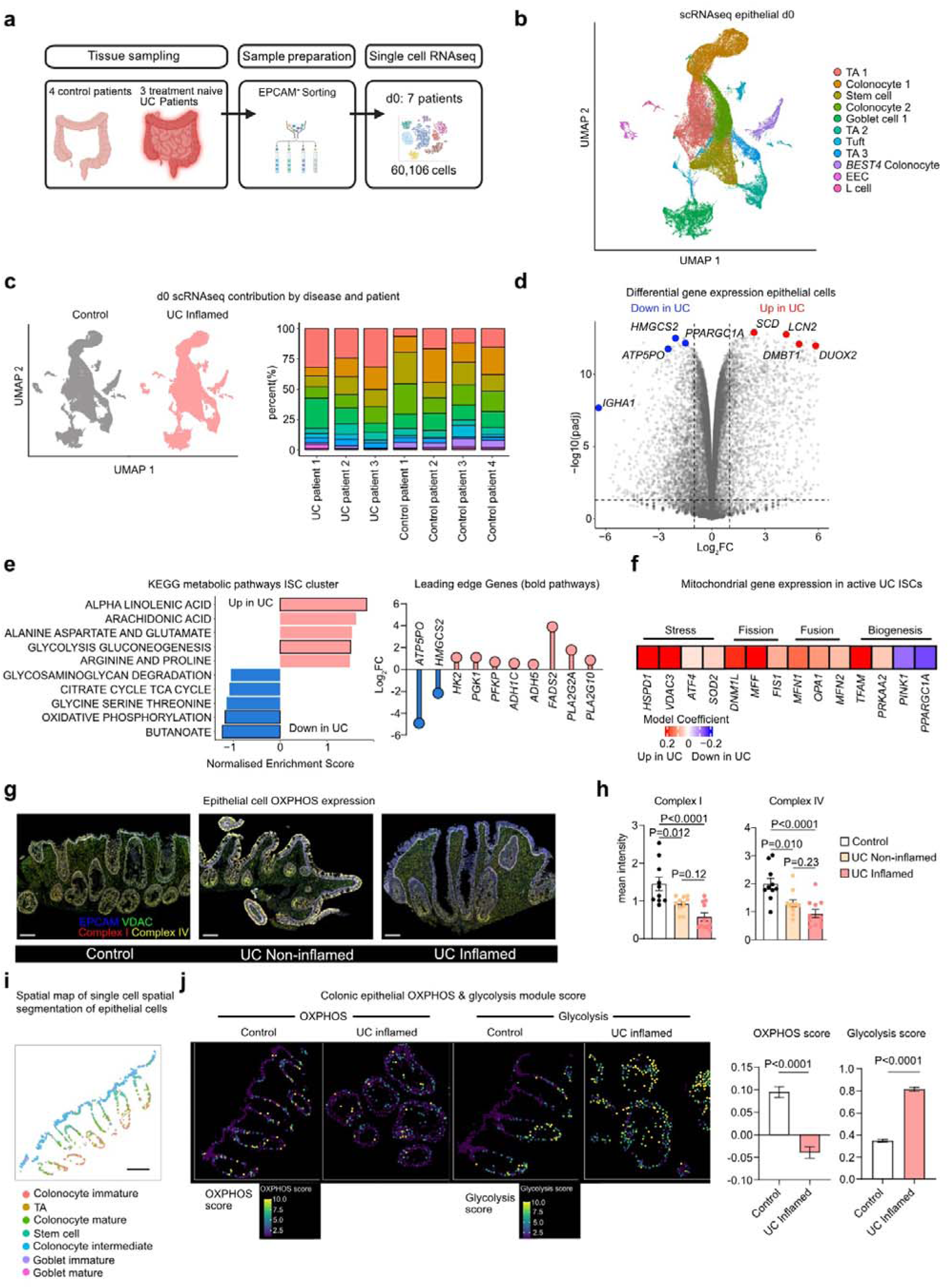
| Metabolic and mitochondrial dysfunction in UC. **a**, Overview of tissue acquisition and processing for scRNAseq. Targeted endoscopic biopsies were acquired from treatment naïve UC patients as well as non-IBD controls. Samples were FACS sorted for EPCAM+. **b**, Unifold manifold approximation and projections (UMAP) dimensionality reduction of epithelial cells from scRNAseq, (n= 4 Control and 3 active UC patients; 27,931 and 32,175 cells, respectively). **c**, Top-UMAP contribution by disease subtype. Bottom-cellular contribution per patient. Legend as per Fig. 1b. **d**, Metabolic changes in epithelial cells in UC. Differential gene expression scRNAseq differential gene expression in epithelial cells (UC active vs control). Top significant genes are highlighted. (pseudobulk via edgR). **e**, Metabolic pathway dysregulation in ISC in UC. Left-shown are changes in normalised enrichment scores of KEGG metabolic pathways UC vs control. Right-Expression of leading edge genes in UC ISCs vs control from bold pathways. **f**, Mitochondrial gene dysregulation in UC. Expression changes (Log2FC, colour bar) of key genes involved in mitochondrial stress and autophagy and biogenesis in inflamed UC ISCs relative to healthy cells. (Black cell outline corresponds to pval adj. <0.05; students t test, Benjamini-Hochberg correction). **g**, Expression of OXPHOS proteins in colonic tissue. Representative immunofluorescence of colonic epithelium across active and non-active UC and matched control patients with staining for EPCAM (blue), VDAC/Mitochondrial mass (green), Complex 1(red) and Complex IV (yellow). (n= 10 control and 11 UC) Scale bar = 100μm. **h**, Normalised expression of Complex I +IV (n=10; one-way ANOVA with Tukey’s multiple comparison test). **i**, Spatial map of single cell spatial transcriptomic segmentation of epithelial cell subtypes in representative field of colonic control tissue, 222,196 cells (157 fields of view (FOV). Scale bar = 100μm. **j**, Switch from oxidative phosphorylation to glycolysis in UC. Shown are quantification of expression of OXPHOS and Glycolysis module score UC inflamed vs control. (n=1/group; 4158 control and 5918 UC; unpaired two-sided Student’s t test).

### Intestinal stem cell mitochondrial and metabolic dysfunction persists in UC organoids

To determine whether metabolic dysfunction persists independently of inflammatory cues—and may thus contribute to the relapsing-remitting course of UC—we analysed UC-derived colonic organoids cultured and differentiated in vitro over 28 days, a time point at which inflammatory conditioning is removed^11^. Following differentiation, we identified relevant colonic cell-lineages including *ITLN1*^+^ *MUC2*^+^ goblet cells, *FABP1*^+^, *MEPA1A*^+^ colonocytes; *PCNA*^+^ (proliferating) and *PROM1*^+^ ISCs (Fig. 2a-c, Extended Fig. 2a and Supplementary Table 4) that are similarly found during earlier time-point at baseline (Day 0), indicating no major shifts in cell populations within the organoid populations at Day 28. We confirmed these differentiated cell types including colonocytes and goblet cells retained enrichment of genes and pathways relevant to their function (Extended Fig. 2b, c). In the absence of local inflammatory cues, genes associated with the emergency epithelial barrier response—such as *REG1A*, *REG1B*, and *PLA2G2A*—were downregulated in UC-derived organoids by Day 28 compared to Day 0 (Extended Fig. 2d). Within ISCs, however, many metabolic genes (*SOD2*, *PPARGC1*α, *ATP5PO*) remained dysregulated, while previously upregulated inflammatory genes (*DMBT1*, *LCN2*, *REG1A*) returned to baseline expression levels (Fig. 2d). Furthermore, similar patterns were observed across the broader epithelial compartment, notably, goblet cells and colonocytes also had similar expression pathways (Extended Fig. 2d, e). To gain further resolution to these transcriptomic changes that were similarly observed in our primary epithelial UC scRNAseq and spatial data (Fig. 1e, i and j), we used flow cytometry metabolic profiling (Single cell energetic metabolism by profiling translation inhibition (SCENITH)) to show functional loss of mitochondrial capacity and heightened glycolysis activity in UC-derived organoids (Fig. 2e, Supplementary Table 5). In addition to this, we also found loss of expression of Complex I and IV proteins (Fig. 2f, Supplementary Table 2f) in UC-derived organoids with consequential slower growth in UC-compared to non-IBD controls (Extended Fig. 2f). Taken together, our data showed a persistence of mitochondrial damage and metabolic dysfunction at in UC ISCs.

**Fig. 2.**
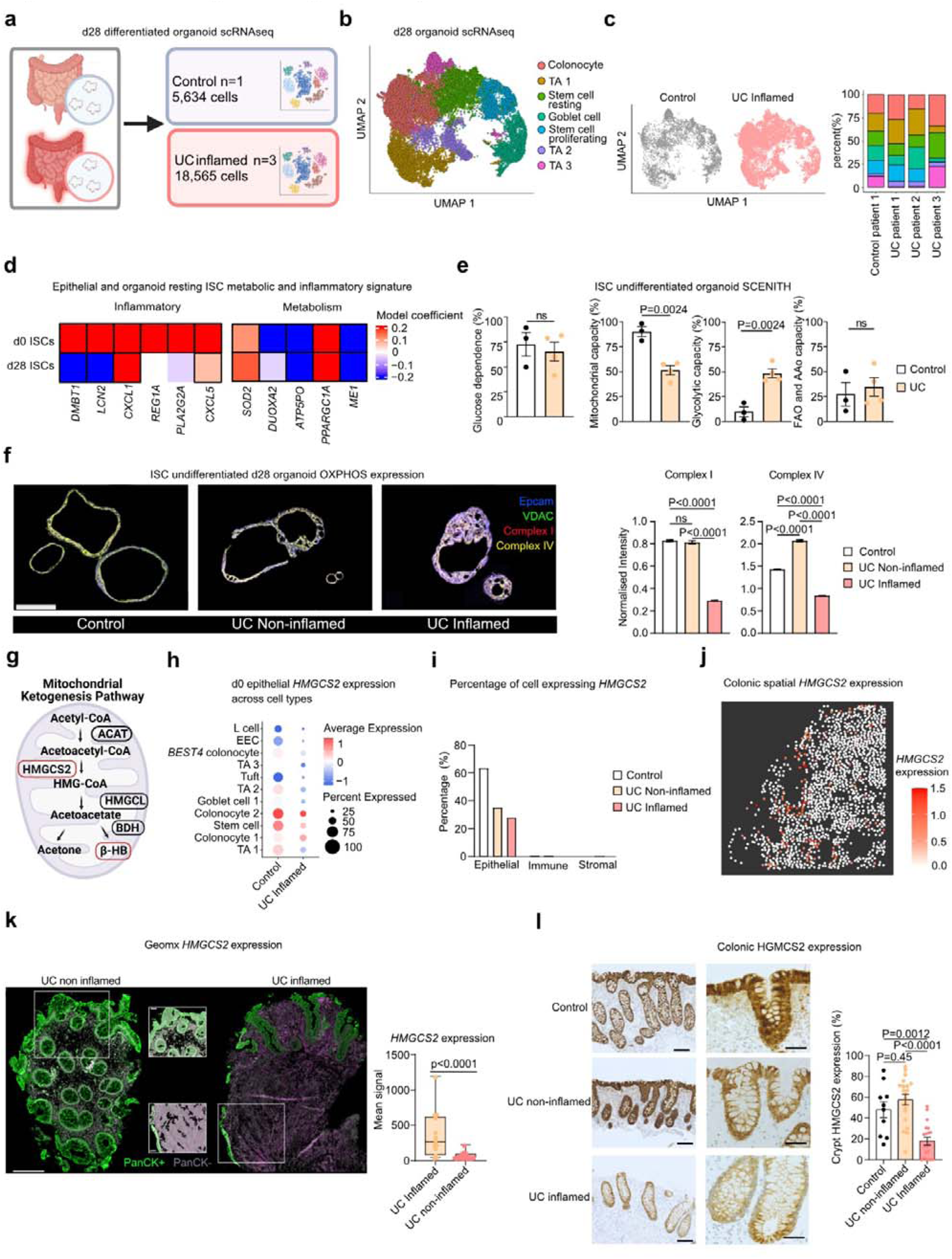
| HMGCS2 gene and protein expression in UC. **a**, Overview of organoid generation for scRNAseq. **b**, UMAP of epithelial organoid scRNAseq (n= 1 control patient (5,634 cells), and 3 UC patients (18,565 cells)) **c**, Top-UMAP contribution by disease subtype. Bottom-cellular contribution per patient. Legend as per Fig. 2b. **d**, Metabolic signature maintained in d28 UC organoids. Expression of key DEG inflammatory and metabolic genes in d0 ISC’s and d28 ISC’s. (UC active vs control, Black cell outline corresponds to pval adj. <0.05, students t test, Benjamini-Hochberg correction). SCENITH (Single Cell Metabolism by Profiling Translation Inhibition) **e**, Bioenergetic classification of undifferentiated organoids via SCENITH, measuring glucose dependence, mitochondrial dependence, glycolytic capacity and fatty acid oxidation (FAO) and amino acid oxidation (AAO) capacity. (n= 3 control 4 UC and, students t test). **f**, Representative immunofluorescence of colonic epithelial ISC enriched undifferentiated organoids across disease states with staining for EPCAM (blue), VDAC/Mitochondrial mass(green), Complex 1(red) and Complex IV (yellow). Scale bar= 100µm. Normalised expression of Complex I +IV across disease state (n=25 organoids/group; one-way ANOVA with Tukey’s multiple comparison test). **g**, Overview of ketogenesis pathway. Fatty acids are transferred to the mitochondria where they undergo β-Oxidation, producing acetyl-CoA units. Two Acetyl-CoA units undergo condensation within the mitochondrial matrix, catalysed by Acetyl-CoA acetyltransferase 1 (ACAT1), producing acetoacetyl-CoA. 3-Hydroxy-3-methylglutaryl-CoA synthase (HMGCS2) then converts acetoacetyl-CoA and another molecule of acetyl-CoA to form HMG-CoA. 3-Hydroxy-3-methylglutaryl-CoA lyase (HMGCL) then cleaves HMG-CoA in acetoacetate and acetyl-CoA. Finally, β-Hydroxybutyrate dehydrogenase (BDH1) converts acetoacetate into β-hydroxybutyrate (βHB). **h**, HMGCS2 expression across epithelial cell types via scRNAseq, control + UC active. **i**, HMGCS2 aggregated expression in epithelial, immune and stromal across control, UC non-inflamed and UC inflamed tissue via scRNAseq. **j**, Spatial transcriptomics displaying HMGCS2 expression across UC colonic crypt. **k**, GEOMX^TM^ spatial transcriptomics platform analysis of HMGCS2 in non-inflamed UC and inflamed UC. Representative images left, scale bar 250 µm, inset 50um. Expression quantified in epithelial panCK+ve region. (n= 15 non-inflamed and n=29 inflamed patients, students *t* test). **l**, Representative images and quantification of colonic epithelial HMGCS2 expression IHC across control and paired, UC non-inflamed and UC inflamed colon (n=10 control, n=20 UC patients; one-way ANOVA with Tukey’s multiple comparison). Results are presented as the mean_J±_JSEM.

### Loss of HMGCS2 function and ketogenesis in UC

In our UC scRNAseq dataset, the most downregulated gene is *3-hydroxy-3-methylglutaryl-CoA synthetase* (*HMGCS2)* (Fig. 1d). *HMGCS2* encodes a mitochondrial-specific, rate-limiting enzyme in ketogenesis, exclusively producing ketone bodies, principally beta-hydroxybutyrate (βHB) (Fig. 2g). Ketogenesis is a conserved cellular response to low energy states, whereby ketone bodies are used as an alternative fuel source^12^. Thus, the finding of *HMGCS2* as the most significantly downregulated gene is surprising in the context of widespread mitochondrial and metabolic dysfunction. In non-IBD colon, *HMGCS2* is highly expressed across all colonic epithelial cells, in particular ISCs and colonocytes (Fig. 2h-j), with significantly higher levels of expression compared to immune and stromal cell populations (Fig. 2i). In UC, we validated the loss of HMGCS2 gene expression using both single-cell and whole-tissue spatial datasets (Fig. 2j–l and Extended Fig. 2 g & h). To gain spatial resolution, in UC, we investigated *HMGCS2* across histological grading of inflammation and found that its expression was lowest in UC mucosa with the most severe inflammation (Fig. 2k). This is confirmed at the protein level, where we showed loss of epithelial HMGCS2 in inflamed mucosal areas of UC as compared to non-inflamed UC and non-IBD controls (Fig. 2l, Supplementary Table 7).

### Ketone body **β**HB restores colonic ISC function in UC

To investigate whether loss of *HMGCS2* expression leads to a functional loss of ketone body production, we measured βHB production *ex vivo* in colonic organoids. Here, we showed significantly lower βHB in organoids from actively inflamed UC-mucosa to non-inflamed UC and non-IBD controls (Fig. 3a, Supplementary Table 8). We next determined if exogenous βHB supplementation could rescue the metabolic shifts we had observed in the UC epithelium and therefore facilitate metabolic re-programming, through its supplementation on D28 differentiated organoids via scRNAseq (Fig. 3b, c, Extended Fig 3a, Supplementary Table 9). All colonic epithelial cell subtypes/populations responded to βHB (Extended Fig. 3b) with upregulation of OXPHOS, TCA and its intermediates (Extended Fig. 3c). In ISCs specifically, βHB elicited a broadly beneficial response, including reduced expression of inflammatory genes (*CXCL1, CXCL5, LCN2, DMBT1*), increased expression of crucial regulators of mitochondrial biogenesis (*PPARGC1*α) and energy production (*ATP5P0*), whilst reducing expression of genes associated with mitochondrial oxidative stress (*SOD2*) (Fig. 3e). Pathway analysis (GSEA) of differentially expressed genes (DEGs) in βHB treated versus vehicle ISCs revealed enrichment related to alternative metabolite metabolism (linoleic acid metabolism), epithelial-immune cross talk signalling (IgA production and toll like receptor signalling) and cell adhesion signalling (Fig. 3f). Next to gain further understanding of the gene regulatory mechanisms that may underpin the preserved metabolic dysregulation in our *ex vivo* organoid systems, we performed transcription factor (TF) enrichment analysis. βHB treatment restored expression of motifs related to secretory cell differentiation (*BHLHA15*), a cellular subtype that is deficient in UC, as well as increased expression of chromatin regulators (*ARID1B, DNMT3B and NCOR1*) (Fig. 3g). Crucially, these chromatin regulators are known to confer epigenetic modulation of *SOX9* essential for ISC homeostasis and regeneration^13^. In support of this, functionally βHB supplementation restored impaired UC organoid growth (Fig. 3h, Supplementary Table 10). In parallel βHB also reduced both cellular and mitochondrial reactive oxygen species, as shown by live confocal imaging of CellROX-stained organoids (Fig. 3i, Supplementary Table 11). These effects were accompanied by improved mitochondrial bioenergetics, evidenced by enhanced mitochondrial capacity and reduced glycolytic activity in βHB-treated UC organoids (Fig. 3j). Taken together, these findings highlight ketogenesis via βHB as a potential metabolic mechanism to enhance epithelial regeneration in UC, possibly through modulation of ISC function.

**Fig. 3.**
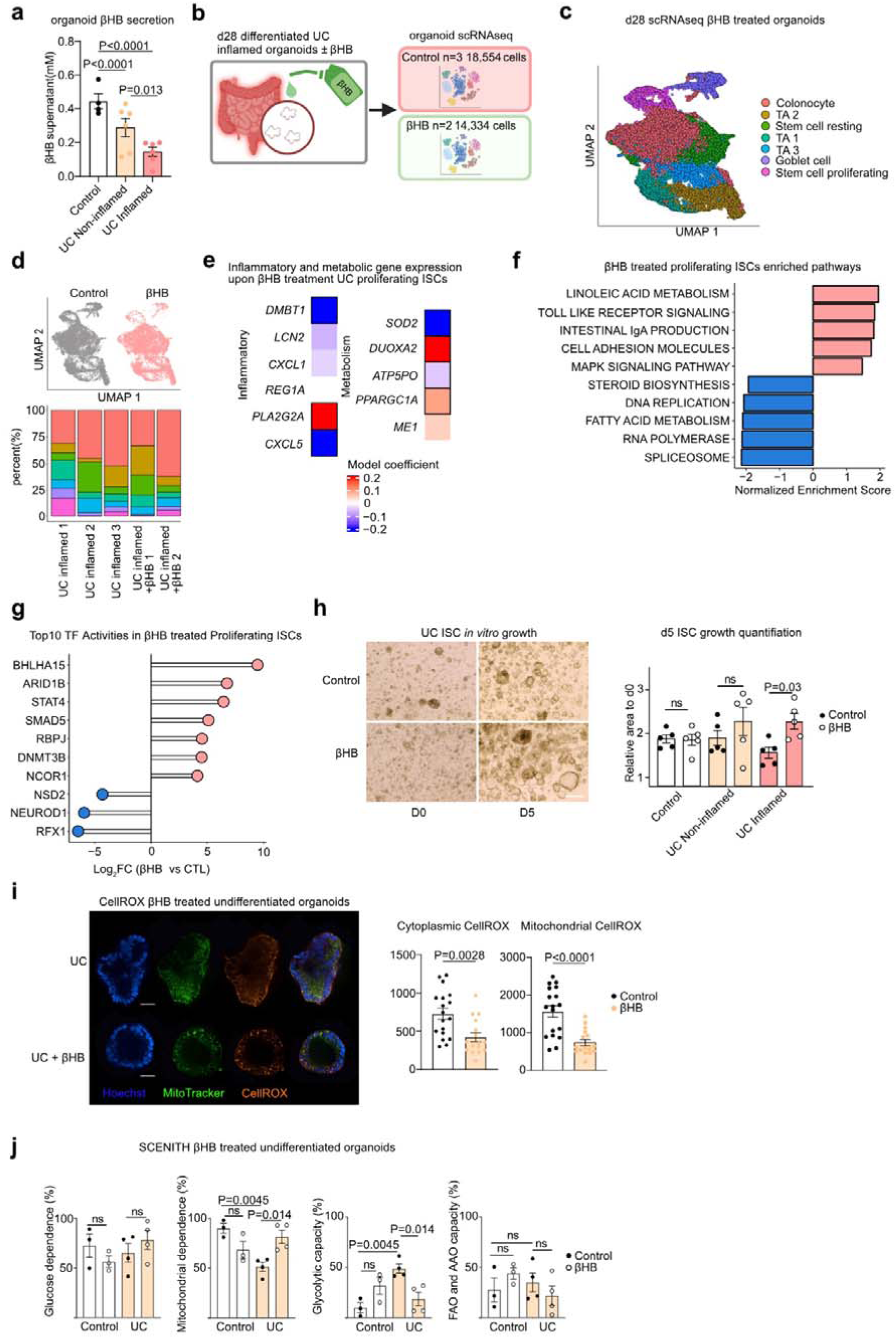
| Ketone body βHB restores colonic ISC function in UC. **a**, Measurement of βHB in organoid media supernatant via colorimetric assay (n=4 control and n=6 UC; one way ANOVA with Sidaks multiple comparison test). **b**, Schematic of βHB treated d28 scRNAseq organoid experiment. **c**, UMAP representation of organoid scRNAseq from UC patients treated with βHB (n=3 UC active, n=2 UC active +0.5mM βHB, 32,000 cells). **d**, Top-UMAP contribution by βHB treatment. Bottom-cellular contribution per patient. Legend as per Fig. 3c. **e**, Changes to metabolic and inflammatory signature upon βHB exposure. Expression of key DEG inflammatory and metabolic genes in βHB vs control proliferating ISC’s. (black cell outline corresponds to pval adj <0.05; students *t* test; Benjamini-Hochberg correction). **f**, Gene set enrichment analysis (GSEA) performed on proliferating ISC βHB treatment vs control, top 10 pathways by normalised enrichment score. **g,** Top variable transcription factors in proliferating ISCs upon βHB treatment (Log_2_FC). **h**, Organoid growth over 5 days in control, UC non-inflamed and UC inflamed populations with and without βHB. Organoids populations were supplemented with 0.5mM βHB and area quantified over 5 days and normalised day 0. (n=5/group; one way ANOVA with SIDAKs multiple comparison test). **i**, Representative image of live confocal imaging of UC organoid ± βHB supplementation for 72hrs *in vitro*, stained with Hoechst (nuclei)-blue, MitoTracker (mitochondria)-green and CellROX (ROS)-orange. Quantification of cytoplasmic and mitochondrial ROS in control and UC organoids with and without βHB. (n=2/group; minimum 16 organoids per group; each dot represents an organoid; students *t* test. **j**, Bioenergetic classification via SCENITH of undifferentiated UC organoid treated with βHB. (n=3/group; one way ANOVA with Sidaks multiple comparison test). Results are presented as the mean_J±_JSEM.

### HMGCS2 in mucosal healing and treatment resistance in UC

Following 28 days of *in vitro* culture, our scRNAseq UC organoid data showed that *HMGCS2* expression remained significantly downregulated (Fig. 4a). Next, we identified hypermethylation at CpG regions associated with regulation of *HMGCS2* in paediatric early-onset UC, suggesting a potential epigenetic mechanism contributing to its reduced expression^14^. To determine whether *HMGCS2* could be therapeutically upregulated, we tested fenofibrate—a known activator of peroxisome proliferator-activated receptor alpha (PPARα), which directly enhances *HMGCS2* expression. Fenofibrate treatment significantly increased *HMGCS2* expression and, importantly, βHB production in UC-derived organoids (Fig. 4c). Next, to assess the translational relevance of these findings, we examined colonic *HMGCS2* expression in UC patients with active disease, with prospective follow-up at six months (www.musicstudy.co.uk). In this longitudinal, paired colonic site-matched sampling, we showed a recovery of epithelial HMGCS2 protein expression that was associated with successful medical therapy and complete mucosal healing at 6 months (Fig. 4d, Supplementary Table 12). Finally, we analysed colonic *HMGCS2* expression from randomised controlled trials of established IBD therapies and found significantly lower *HMGCS2* expression with treatment resistance to four anti-inflammatory biologic therapies (Fig. 4e), including anti-TNF (infliximab and golimumab), anti-IL-12/23 (ustekinumab) and anti-α4β7 (vedolizumab). Taken together, these lines of data suggest that HMGCS2 can indeed be pharmacologically targeted, and given its association with clinical outcomes, may prove an attractive therapeutic target with wider improvements in mitochondrial and metabolic colonic epithelial function in UC (Fig. 4f).

**Fig. 4.**
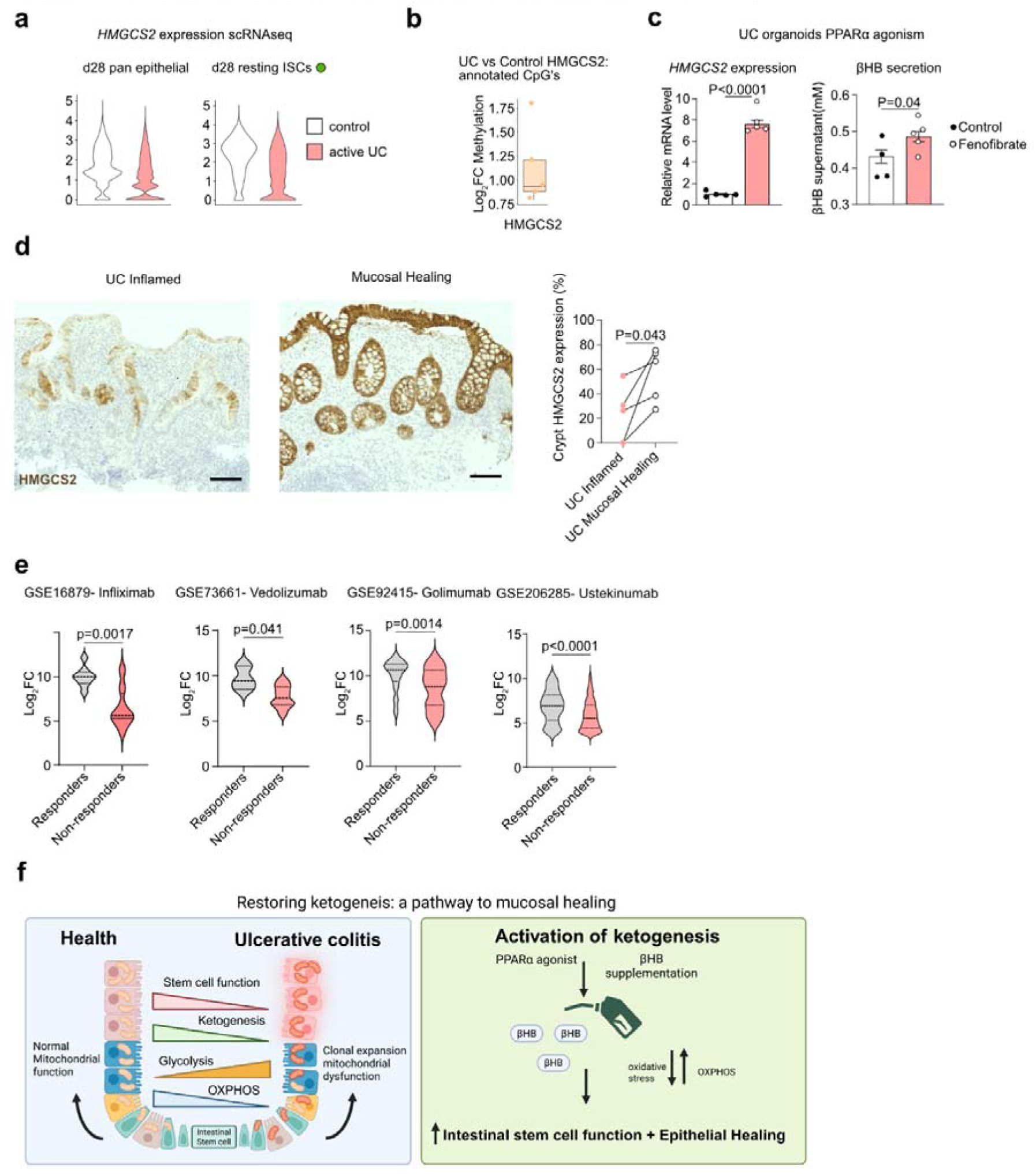
| HMGCS2 and resolution of inflammation in UC. **a**, *HMGCS2* expression via scRNAseq of all epithelial cells and in d28 organoid derived ISCs. Pseudobulk via edgR. **b**, Expression of *HMGCS2* annotated CpG’s in paediatric UC vs control colon (n= 24 control and 18 UC) **c**, PPARα agonist treatment of UC organoids with fenofibrate (25µm), 48hrs *in vitro* culture. Left, *HMGCS2* expression in UC organoids (n=5/group) by qPCR. (n= 5 independent pools of organoids from 1 donor; students *t* test). Right, βHB in organoid media supernatant via colorimetric assay (n=5/ group; students *t* test). ***d***, HMGCS2 expression by IHC, across inflamed UC and sites that subsequently responded to medical therapy displaying mucosal healing (scale bar 100µm). (n= 4 paired patients; paired students *t* test) **e**, *HMGCS2* expression in responders and non-responders to medical therapy (left to right, Infliximab, Vedolizumab, Golimumab, Ustekinumab) from colonic microarray (GSE1689 n=8 responders, 16 non-responders; GSE73661 n=3 responders, 21 non-responders; GSE92415 n=39 responders, 36 non-responders; GSE206285 n=95 responders and 451 non-responders); Mann-Whitney U test). **f**, Schematic of the role of ketogenesis as the tipping point for metabolic failure in UC ISCs. Through restoration of ketogenesis in UC ISCs cellular stress is reduced, OXPHOS capacity is enhanced and ISC function is enhanced. *HMGCS2* activity is strongly linked to mucosal healing in UC, potentially revealing ketogenesis as a novel approach to restore colonic ISC metabolic dysfunction in UC. Data in a-d are mean_J±_JSEM, data in e are median ± IQR.

## Discussion

We present multiple levels of evidence to support a disease-specific mechanism implicating a failure of ketogenesis at the ISC level in UC. A colonic epithelial ‘metabolic block’ in UC promulgated by microbial dysbiosis, mitochondrial injury and genetic predisposition has long been described. In the original ‘energy-deficient’ hypothesis, Roediger et al. described a failure in mitochondrial fatty acid oxidation and interestingly, also ketogenesis^15^. In UC, our data points to hitherto unknown loss of HMGCS2 function within ISCs, the rate-limiting enzyme to produce ketone bodies acting as ‘back-up energy’ failure and, this is potentially the tipping point towards a pathogenic point of epithelial metabolic dysfunction with its effect imprinted within the ISCs (Fig. 4f).

HMGCS2 is highly expressed in the colonic epithelium indicating the importance of this metabolic pathway where locally produced ketone bodies (previously thought to be hepatocyte specific) are vital for ISC function^16^. Recent studies have demonstrated low *HMCGS2* expression in UC epithelial cells in support of our findings^17–19^. Furthermore, in the mouse, conditionally deletion of *HMGCS2* within the intestinal epithelium resulted in defective ISC function with increased susceptibility to radiation induced colitis ^16^. More specifically, in an alternative intestinal *HMGCS2* knock out model mitochondrial metabolism was specifically affected with reduced expression of OXPHOS pathways and breakdown of epithelial barrier function^20^. In our study, we provide crucial evidence of loss of ketogenesis via HMGCS2 in UC patients, demonstrate that this pathway is pivotal to ISC function in the human epithelium and link expression of HMGCS2 to clinical outcomes.

Whilst data we present demonstrate that *HMGCS2* is hypermethylated in the UC epithelium, the precise upstream mechanisms and regulatory factors driving the loss of *HMGCS2* expression remain undefined. It is possible that impaired ketogenesis is a secondary consequence of extensive mitochondrial damage in UC-derived colonocyte populations. Moreover, we do not differentiate whether the effects on ISC function arise directly from energy deficiency or from broader signalling roles of ketone bodies. For instance, β-hydroxybutyrate (βHB) has been shown to inhibit the NLRP3 inflammasome and act as a histone deacetylase (HDAC) inhibitor^16,21^. Finally, the absence of IBD phenotypes in individuals with inherited mitochondrial disorders suggests that primary defects in cellular energetics alone are insufficient to drive colitis, and/or that compensatory pathways may exist to mitigate against bioenergetic stress in the gut^22^.

UC is a difficult condition to treat. Despite immune- and anti-inflammatory treatments, disease relapses and flares are very common with a significant proportion of UC patients failing to achieve complete mucosal healing^23^. Our findings, therefore, have direct translational implications. In mouse, both ketone diets and βHB topical therapy have been shown to improve induced colitis^24,25^. Our work opens up a simple and novel therapeutic approach in UC to promote mucosal healing and remission via a metabolic approach. This can be achieved by pharmacologic targeting of HMGCS2 within the colon epithelium; or more indirectly, by inducing ketogenesis by diet or fasting. In other metabolic conditions such as metabolic dysfunction-associated steatohepatitis (MASLD), diabetes and heart failure, promoting ketogenesis appeared to have a beneficial effect^26–28^. Hence, our findings offer a new therapeutic dimension that complements current advanced anti-inflammatory therapies. In this wholly human UC study, we provide multiple levels of evidence to implicate ketogenesis as a tractable target and thereby the credence to pursue this translational angle.

## Materials and methods

### Human subjects

The acquisition of human samples was approved by National Health Service (NHS) Research Ethics Committee (REC 18/ES/0090) and (REC 19/ES/0087) as part of the GI-DAMPS (Investigation into Gastrointestinal Damage Associated Molecular Patterns) and MUSIC (Mitochondrial DAMPs as mechanistic biomarkers of mucosal inflammation in Crohn’s Disease, www.musicstudy.uk) studies. Written informed consent was given by all participants. Individuals undergoing routine endoscopic assessment were recruited at the Western General Hospital, Edinburgh, UK. All UC participants had macroscopic inflammation and a confirmed histological diagnosis. Endoscopic assessment of disease severity at the biopsy site was performed by clinicians using the Mayo endoscopic sub-score for UC. All control participants were undergoing endoscopic assessment for non-cancer related reasons, with macroscopic and histological confirmation of normal colonic tissue. For scRNAseq and organoid generation 4-6 biopsies of were taken and deposited into pre-chilled DMEM+++(Advanced DMEM/F-12 /1%Hepes/2mM GlutaMAX (all Gibco)/10%FCS (Sigma)/100µg/mL Primocin (InvivoGen). Samples were placed on ice and immediately transferred for processing. For histological analysis of samples, biopsies were immediately placed in 4% paraformaldehyde (PFA).

### Sample information

Patient characteristics for each experiment are supplied in Supplementary Tables 1-12.

#### Single cell dissociation from fresh biopsies

Single-cell suspension from collected biopsies (4-6) were obtained using a modified version of a previously published protocol utilizing and enzymatic digestion and mechanical dissociation technique^29^. All pipette tips were coated in a 4°C 1% BSA (Sigma)/DPBS (Ca/MG-free) solution. Biopsies were washed x3 in 1% Penicillin-streptomycin (Gibco) in DBPS. Then incubated in a shaking incubator (150rpm) twice for 10 min at 37 °C in R5 (5mls RPMI/5%FCS/1%PenStrep) containing 4mM DTT (Medkoo). Biopsies were then incubated in a shaking incubator (150rpm) in 5ml HBSS/5mM EDTA (Gibco) for 10 min at 37°C. Biopsies were then digested to single cell suspension by incubation in a shaking incubator (200rpm), at 37°C in 5ml R5 containing 30µg/mL Dnase I (Roche) and 0.2mg/mL Liberase TM (Roche). Digestion was monitored for each sample and once whole tissue visibly digested (typically 30mins) the enzymes were inactivated with 30ml of 4°C DMEM++++. Single cell suspension was passed through 70µm cell strainer, resuspended in 1mL of Red cell lysis solution (Roche) for 1 min at room temperature and washed twice more in DMEM++++.

#### Flow cytometry assisted cytometric *sorting (FACS)* of colon tissue

Intestinal epithelial cells were purified by FACS, thus: Single cell suspensions of colonic tissue resuspended in 200µl of FACS buffer(DPBS(Ca/MG-free), 2% FCS, 2mM EDTA) and stained with PE anti-CD326 (EPCAM) (324206, Biolegend,1:100) BUV 393 anti-CD45 (563791, BD biosciences, 1:100), APC anti-CD-90 (328114, Biolegend, 1:200) and BUV 480 anti-CD31 (566144, BD biosciences, 1:25) for 1 hr at 4°C, Sytox BB515 (S7020, Invitrogen, 1:8000) was added for 10mins before acquisition to exclude dead cells. Live CD34^-^, CD90^-^, EPCAM+ cells were purified by FACS on a BD Aria Fusion 6 (BD Biosciences). Purified epithelial cells were fixed for subsequent library preparation as per manufactures instruction (CG000478, Rev B). Samples were stored at −80°C for contemporary batch preparation of libraries.

### scRNAseq

#### Probe based scRNAseq library preparation and sequencing

Both d0 and d28 organoid epithelial samples were processed using the protocol according to manufacturer’s instructions (CG000478, rev D). Cells were counted on LUNA-FX7 cell counter (AO/PI viability kit, Logos), targeting 12,000 cells per sample. Single-cell suspensions were converted to barcoded scRNAseq library using the Chromium 3’ reagent kits v3.1 (10x genomics), as per manufacture’s protocol for Chromium Fixed RNA profiling for multiplexed samples (CG000527, rev E). Libraries were sequenced using a NovaSeq 2000 (100bp)/ X (150bp) sequencer aiming for 70,000 read pairs/cells according to manufacturer’s protocol. Bcl2fastq was used for demultiplexing libraries after sequencing. FASTQ files were processed with Cell Ranger 7.1.0 (10X Genomics) with multi pipeline and human genome reference GRCh38-2020-A.

#### Data processing

Processing of scRNAseq data was conducted identically for each of the four scRNAseq datasets presented. The raw feature barcode matrix was processed in R-Studio (R version 4.3.1), using Seurat (version Seurat v5.0.1)^30^. Cells with > 200 detected genes and < 20% mitochondrial reads were retained. To control for batch effects, the filtered count matrices were normalized using SCTtransform, regressing on mitochondrial and ribosomal reads with the method set to ‘glmGamPoi’^31^. Doublets were identified using DoubletFinder, expecting 5% doublets, with principal components 1-10, pN = 0.25, and pK = 0.09^32^. Next, the Seurat function SelectIntegrationFeatures was used with nfeatures set to 3000. This was followed by principal component analysis (PCA) using RunPCA with npcs = 30. The data were then integrated using the IntegrateData function, utilizing reciprocal PCA integration (RPCA) based on SCT-transformed data. Next, the FindNeighbors function was applied, using the first 30 PCA dimensions as input features. Clustering was performed with the FindClusters function with the expression level of canonical marker genes and the top differentially expressed genes were used for identifying known cell types corresponding to the clusters^2,17,33^. A resolution of 0.15 for primary epithelial cells and 0.15 for organoid cells was used. Finally, dimensionality reduction was conducted via RunUMAP. Total analysed cells per dataset after QC: d0=63,634, d28= 24,199, d28+βHB= 32,888, publicly available dataset= 308,442^17^.

#### scRNAseq analysis

Differential gene expression analysis was performed using pseudobulk analysis (EdgR) via DElegate^34^. Pathway analysis was conducted using fgsea (version 1.23.0)^35^. Transcription factor inference was conducted using decoupleR (version 2.6.0)^36^. Gene enrichment analysis was conducted by conducting an ordinary least squares linear regression model, with statistical testing corrected using the Benjamini-Hochberg FDR method^37^. Heatmaps were plotting using R package ComplexHeatmap (version 2.16.0).

### Human organoid culture

#### Cell isolation and organoid culture

Human organoids were generated from digested single cell suspension of colonic biopsies described prior. Cell pellet was resuspended in 15µL of Matrigel (Corning) per biopsy. Matrigel was dispensed onto a pre-warmed 24 well culture plate, inverted and upon solidification of Matrigel droplets overlayed with the following media ^38^: LWRN conditioned media 50% (manufactured as described by Dr. Thaddeus Stappenbeck’s Lab^39^, DMEM adv/f12, 10 mM HEPES, 2 mM Glutamax, B27 (1x) (all from Gibco), 1 mM N-acetylcysteine (Sigma), 50 ng/mL epidermal growth factor, 10 mM Nicotinamide (Sigma), 500nM A83-01 (Bio-techne), 10nM Prostaglandin (Bio-techne), 10 nM gastrin (Sigma), 10 µM SB202190 (Sigma), 2.5 µM Thiazovinin (Stratech), 2.5 µM CHIR99021(Bio-techne), 10 mM Y-27632 (Bio-techne), 100µg/ml Primocin. Following establishment of organoids (48-72hrs) Y-27632 dihydrochloride and CHIR99021 were removed from media. Following this 500µL, initiation media was added and changed 3x weekly. Differentiation was achieved through reduction of conditioned media concentration to 25% for 5 days and further 2 days at 12.5%, alongside removal of EGF, Nicotinamide, A83-01, Prostaglandin, SB202190, Thiazovinin and addition of 100ng/ml IGF and 50ng FGF/ml (both Peprotech)^40^. Where applicable organoid cultures were supplemented with 25µm Fenofibrate (10005368, Cayman) and 0.5mM Sodium β-hydroxybutyrate (54965, Sigma) for 48hrs.

#### Recovery of organoids from 3d growth matrix

Organoids were released from Matrigel with cell recovery solution (Corning) at 4°C. Organoids were centrifuged at 450g for 5mins and supernatant and Matrigel aspirated. If there was residual Matrigel was present a further wash in cell recovery medium was performed for 10mins.

#### RNA extraction from organoids

After isolation of organoids from Matrigel, pellets were extracted by using the TRIzol Purelink RNA mini Kit (12183018A, Invitrogen). The RNA was quantified using Nanodrop (ThermoFisher Scientific), mRNA was reverse transcribed to cDNA (High-Capacity Reverse transcription kit; ThermoFisher Scientific). Gene expression analysis was performed using the standard curve method with specific primers to *GAPDH (Forward-* AAGGTGAAGGTCGGAGTCAAC, Reverse-GGGGTCATTGATGGCAACAATA) and *HMGCS2 (Forward-* TACCACCAATGCCTGCTACG, Reverse-TGGCATAACGACCATCCCAG) using SYBRGreen (Applied Biosciences) with the QuantStudio 5 Real-Time PCR System (ThermoFisher Scientific).

#### Size analysis of organoids during *in vitro* culture

Organoid size was quantified from 24hrs post passage for 5 days. A minimum of 3 brightfield fields of view were captured per technical replicate and timepoint, using an EVOS XL core microscope (ThermoFisher Scientific). Organoid images were analysed using OrganoID, with area at d5 normalised to d0^41^. Areas of a minimum of 100 organoids per timepoint were quantified.

#### Preparation of organoids for probe based scRNAseq

Organoids were isolated from Matrigel as above. 500ul of 37°C TrypLe (Gibco) was added, dissociation being monitored closely under a microscope. 10ml ice cold DMEM++++ was added and cells were centrifuged at 500g, 4°C for 5mins. Cells were strained using 70µm cell strainer and dead cells were removed using the EasySep dead cell removal (Annexin V) kit (17899, Stemcell Technologies). Isolated cells were then processed as for d0 epithelial cells using 10x protocol described above.

#### Generation of organoid sections

Organoids were isolated from Matrigel as before. Organoids were fixed for 20 mins at 4°C in 4% PFA^42^. Post fixation organoids were washed twice in DPBS and then were resuspended in 100µl 1% low melting point agarose (ThermoFisher Scientific), placed in 7mm mould (CellPath), and allowed to solidify for 1 hr at 4°C. Agarose blocks were transferred to a 70% ethanol solution for storage, and sectioned to 4µm thickness.

### Single Cell Energetic metabolism by Translation (SCENITH)

Organoids were disassociated to single cells as above and SCENITH conduced as per Argüello et. al. ^43^. Briefly, organoid pellet for each sample was resuspended in DMEM++++ at 37°C and incubated for 30 min at 37 °C, 5% CO2. Biological replicates were tested in triplicate. Samples were then treated for 15 min at 37 °C, 5% CO2 with control, 2-DG (Cayman, 100mM), oligomycin (Cayman, 1µM), harringtonine (Cayman, 2µg/ml) or a combination of both oligomycin and 2-DG (2-DG for 10mins, then oligomycin added). Puromycin (ThermoFisher Scientific: 10µg/ml) was added for further 30mins at 37 °C, 5% CO2. Samples were then washed x2 in FACS buffer. Samples were then resuspended in Fc Block (422302, Biolegend,1:200), and incubated in dark for 10 min at 4°C, followed by addition of fixed viability dye (65-0863-14, eBioscience, 1:400) for further 10min. Samples were washed in FACS buffer and then fixed and permeabilised using the Foxp3 fixation/ permeabilization kit (00-5521-00, eBioscience), following the manufacturer’s instructions. Samples were resuspended in intracellular block (1x perm buffer +20% FCS) at room temperature for 10 min, followed by addition of anti-puromycin antibody (381508, Biolegend, 1:500) for 60mins at 4°C. Samples were washed, cell strained (40µm) and resuspended in 200µl FACS buffer. Analytical flow cytometry was performed using a Cytek Aurora cytometer (Cytek) and data analysis was performed using FlowJo (version 10.10.0).

### Immunohistochemisty and Immunofluorescence

#### Immunohistochemistry

Colonoscopic biopsies were fixed in 4% PFA and sectioned at 4µm. Sections were de-paraffinized and rehydrated. Antigen retrieval was performed via pressure cooking in Tris 1mM EDTA Ph 9 buffer for 5 min. Samples were blocked in 3% hydrogen peroxide (H1009, Sigma) solution for 20 min at room temperature, followed by 2x wash in PBS Triton 0.1% (X100, Sigma). Blocking was performed with 2.5% normal horse serum (MP-7401, Vector laboratories) for 1hr at room temperature. Sections were then incubated in anti-HMGCS2 antibody (ab137043, Abcam,1:500) for 16hr in a humidified chamber at 4°C, followed by detection via secondary antibody: Horse anti-Rabbit IgG (MP-7401, Vector) for 30 mins at room temperature. Detection was performed using the ImmPACT DAB EqV kit (SK-4103, Vector) as per manufactures instructions. Non primary controls were performed for each experiment. Slides were imaged with Axioscan Slide Scanner (Zeiss). Scanned images were analysed in a blinded fashion using QuPath v0.4.2^44^. Haematoxylin and DAB stain vectors were set according to a positive and negative control. A minimum of 3 sagittal crypts per slide were manually annotated per slide, and DAB positive area was calculated.

#### Immunofluorescence

Antigen retrieval was performed via pressure cooking in 1mM EDTA buffer ph 8.0. Non-specific binding was blocked in 10% normal goat serum (S-1000, Vector). Followed by avidin/biotin blocking, as per manufactures instructions (SP-2001, Vector). The primary antibodies EPCAM (ab124825, Abcam, 1:80), Anti-NDUFB8 (IgG1) (ab10242, Abcam, 1:40), Anti-MTCO1 (IgG2a)(ab14705, Abcam, 1:50), anti-VDAC (IgG2b)(ab14734, Abcam, 1:100) were added and incubated overnight in a humidified chamber at 4°C. Detection was performed with addition of Alexa Fluor 405(1:150), Anti-IgG2b 488 (1:200), Anti-IgG2a 546(1:200), Anti-IgG1 biotin(1:200) anti-goat secondary antibodies (all Invitrogen) for 2hr at 4°C and further incubated with streptavidin conjugated with Alex 647 (Invitrogen, 1:200) for a further 2hr at 4°C. Sections were washed and mounted in Prolong gold (Sigma). Non-primary antibody controls were processed for each sample. Organoid sections were derived from organoids cultured over at 6 weeks and were processed for immunofluorescence in an identical manner.

#### OXPHOS protein quantification

Stained sections were imaged using a Sp8 confocal microscope (Leica microsystems) and analysed using Imaris software (Bitplane, version 9.6). Briefly, epithelial cells were identified and mean optical density of each cell was calculated with correction for non-primary control.

### Live organoid immunostaining

Organoids were recovered from matrigel as above. Whole organoids were resuspended in 300ul DMEM+++ at 37°C for 30 min, with addition of 5µM CellROX far red (C10422, Invitrogen) and 200nm of MitoTracker Green (M7514, Cell Signalling) 20µM Hoescht (33342, Thermofisher Scientific) was added for final 10 min. Organoids were washed in DMEM+++ at 37°C twice and then resuspended in 10µl 1% low melt agarose (37°C), which was then plated in an µ-Slide Angiogenesis IbiTreat slide(81596, Ibidi). The slide was pulse centrifuged for 30sec at 100g at 4°C, followed by incubation at 4°C for 2 mins before addition of 50ul DMEM+++/well. Samples were then imaged immediately.

Imaging was performed on Opera Phenix Plus (Revvity). A minimum of 9 organoids were imaged using 63x objective (NA 1.15) for each biological replicate. Image analysis was performed using Revvity’s Harmony analysis software. Cell nuclei were identified via Hoechst staining, and mitochondrial puncta identified via MitoTracker. A minimum of 50 nuclei and 200 mitochondria were captured per organoid. Intensity of CellROX staining within cytoplasm and individual mitochondria was calculated, after subtraction of background fluorescence. Following this the average CellROX intensity was calculated for each organoid.

### Colorimetrics

Extracellular βHB was measured by sampling organoid media from organoids that were isolated from paired inflamed and adjacent non-inflamed colon, using a colorimetric assay kit (ab83390, Abcam) in technical triplicate using the Biotek Synergy HT plate reader (Agilent). 50µl of organoid media was sampled after 48hrs of *in vitro* culture, media was diluted 1:2 in assay buffer and protocol performed as per manufactures instructions. Absorbance was measured using Biotek Synergy HT plate reader.

### Spatial transcriptomics

#### GeoMx whole transcriptome digital spatial profiling

Endoscopic punch biopsies from colitis patients from the UNIFI study^45^ (double-blind, placebo controlled trial) were collected prior to treatment, fixed in formalin and embedded in paraffin blocks for archiving (NCT02407236). Blocks from 45 subjects were selected at either extreme of the Geboes scale (n=29 for score 15-20, n=16 for score 0-4). Slides were processed for digital spatial profiling on the GeoMx platform under basic antigen retrieval conditions and epitope unmasking as previously described^46^. Briefly, the whole transcriptome atlas panel was incubated ∼20hr for hybridization, followed by stringency washes to remove nonspecific binding, and finally staining for reference morphology markers (Syto13, PanCk, CD45; Bruker Spatial, Seattle). Slides were imaged on the GeoMx instrument and regions of interest (ROI) of 500μm by 500μm were placed. Patients ranged from 2-9 ROIs per tissue. Individual ROIs were segmented based on morphology staining into two areas of illumination (AOI): PanCk+ and PanCk-nucleated cells. Mean of each ROI per patient were calculated per segment and compared.

### CosMx spatial molecular imaging (SMI)

#### 6000plex

Endoscopic punch biopsies from untreated colitis patients (n=4) were processed for single cell profiling by spatial molecular imaging on the CosMx platform (Bruker Spatial, Seattle) measuring the expression of 6180 genes (40 FOV/sample). Initial segmentation was performed using CellPose^47^. Cell-by-gene count matrices and cell-centroid data were further filtered to the threshold minimum of 60 transcripts per cell, yielding 208,956 cells. Next, gene count data was log-normalized (Seurat v5) and label transfer was performed with SingleR^48^. Public IBD scRNAseq data was used as a reference for cell type annotation^17^. Highly variable genes (HVGs) were computed using modelGeneVar^49^ and intersected them with the gene features present in the spatial dataset. For the initial low-resolution label transfer, the top 100 upregulated DEGs for each major cell type (epithelial, myeloid, B/plasma, stromal, and T/NK) from the intersected HVGs using findMarkers. These top DEGs, reference scRNAseq, spatial query, and major cell type labels of reference were used as inputs for the label transfer with SingleR. To refine annotations beyond the six broad categories, we divided both the reference and spatial query data into the five major cell type compartments and repeated the label transfer. In this step, we followed the same procedure but used only the top 40 DEGs per granular cell type.

#### 1000plex

We used the Nanostring CosMx Spatial Molecular Imaging platform to measure expression of 960 genes discriminating transcriptional profiles and spatial localization of 70,731(87 FOV) and 151,465(70 FOV) cells from 1 Control and 1 UC active colon, respectively. Initial image segmentation was performed with Mesmer ^50^ with the following parameters: mesmer_mode = “both”, scale = pixel size of the images. Subsequently, initial segmentation boundaries were used as a basis for refinement with Baysor^51^. In Seurat (v5.3.0), quality control was performed by filtering out cells with < 25 counts, followed by log-normalisation. Cells that passed this filtering step were log-normalised to mean counts and scaled using Seurat (v5.3.0). Principal component analysis (PCA) was conducted and used to generate a two-dimensional UMAP. Cells were clustered using a k-nearest neighbours’ approach (k.param) based on the PCA space. Broad cell types - immune, stromal, and epithelial - were annotated based on DEG markers and PanCK, CD45, and CD68 labelling. The epithelial cell cluster was subsetted and normalised and scaled in followed by integration using Harmony (v1.2.3)^52^. Post-integration, cells were re-clustered and annotated using a semi-supervised approach, incorporating reference expression data from the Gut Human Cell Atlas. Cell boundaries were inferred and visualised using the CellPoly package (v0.0.0.9). Glycolysis and oxidative phosphorylation (OXPHOS) module scores were computed and visualised using CosMx gene sets.

### Analysis of publicly available scRNAseq, microarray and DNA methylation data

The following datasets were accessed from the Gene Expression Omnibus: (1) scRNAseq data of immune, stromal and epithelial cell compartments of UC patients Single Cell Portal: SCP259^17^; (2) normalised genome-wide DNA methylation data of paediatric UC patients sequenced on the Illumina Infinium Human Methylation 450 and EPIC BeadChip platforms, acquisition: E-MTAB-5463^14^; (3) microarray data from colon biopsies of responders and non-responders to TNF blockade, anti-α4β7-integrin GSE73661^53^, anti-TNFα GSE73661^54^, anti-TNFα GSE92415 ^55^ and anti-IL-12/23 GSE206285^56^.

#### CPG analysis

Differentially methylated CpG probe data were obtained from a published dataset (Howell et al., 2018; Supplementary Table, mmc12.xlsx). CpG probe identifiers were matched to gene annotations using the IlluminaHumanMethylation450kanno.ilmn12.hg19 R package via the getAnnotation() function from the minfi package (version 1.46.0)^57^. Where multiple genes were annotated to a single probe, the first listed gene was retained for downstream analysis. Genes of interest, such as HMGCS2, were further interrogated by extracting all annotated CpG probes and visualizing their methylation changes using boxplots of logFC values.

### Statistical analysis

Statistical details of experiments can be found in the figure legends and the results section including the sample number, information on definition of centre and dispersion and type of statistical tests used to associate or compare groups. Statistics and graph plotting were performed using Prism 9 (GraphPad Software).

#### Figure artwork

Figure artwork was created with BioRender.com.

## Acknowledgements

We are grateful to Dr Justyna Cholewa-Waclaw, IRR Imaging Centre, for performing live confocal organoid imaging. Flow cytometry data were generated with the support from the IRR Flow Cytometry and Cell Sorting Facility. We would also like to thank Meryam Beniazza from the IRR single cell facility for performing scRNAseq library preparations. Furthermore, we would like to thank Melanie McMillan for her expertise in CosMx slide preparation. Finally, we gratefully acknowledge the Newcastle University BioImaging Unit for their support & assistance in this work.

## Author Contributions

G.T.H. conceptualized the study. V.E.C., R.W., A.J.B.C., P.D.C., P.Y.L., M.H.I.B., C.E.A., C.D.K, D.R., J.R., K.H., A.S., and G.R.J performed and analysed experiments. L.C.G. and N.B.J. edited the manuscript. D.R. wrote the original draft of the manuscript. G.R.J. and G.T.H. supervised the study.

## Data and material availability

Data supporting the findings of this study are available within the article and Supplementary Information and from the corresponding author on reasonable request.

---

**Extended Data Fig. 1.**
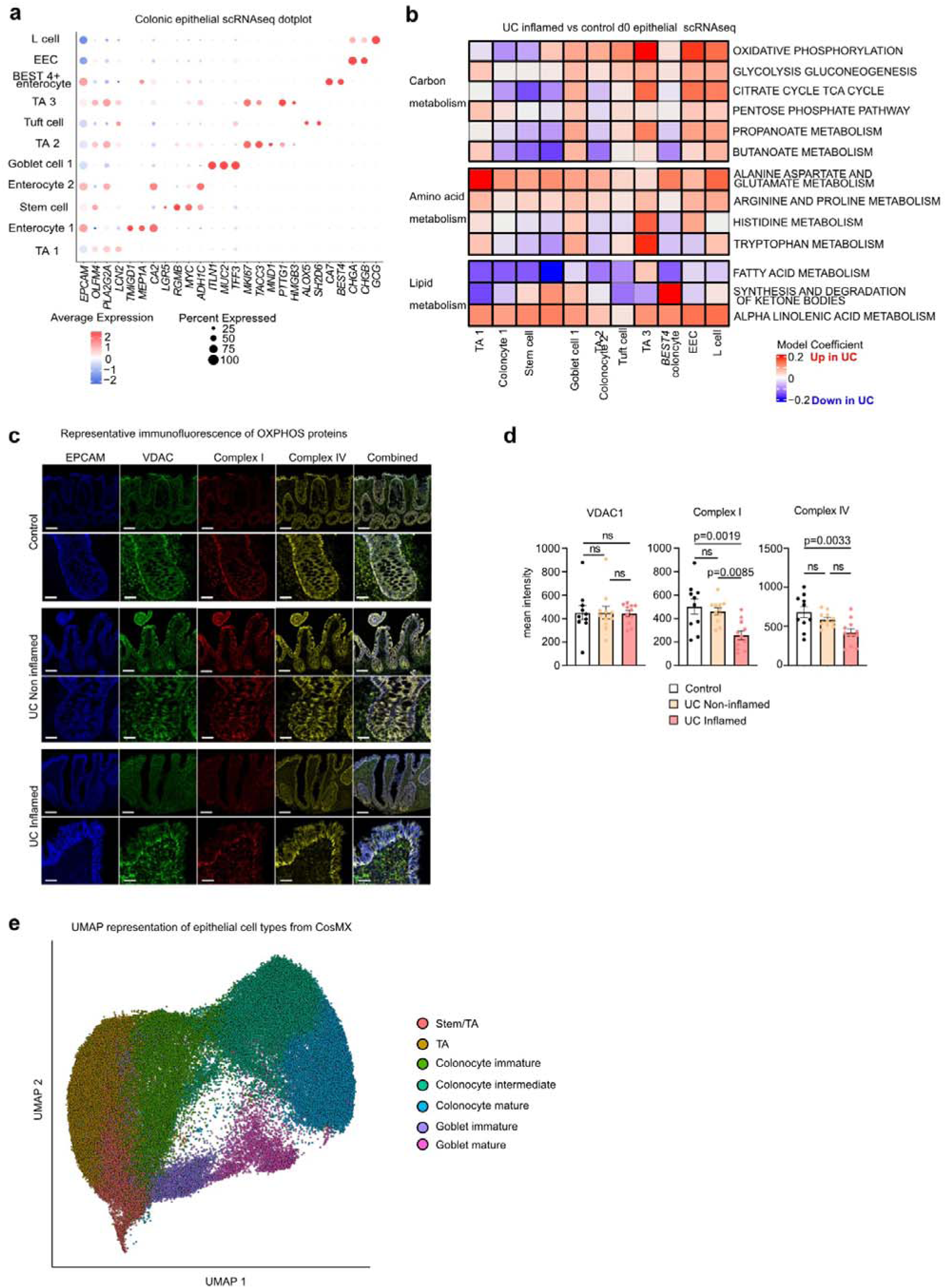
| Dysregulation of Oxidative phosphorylation in UC epithelium. **a,** Expression of key marker genes for each scRNAseq d0 epithelial cell type. **b**, Metabolic dysregulation across epithelial cells in UC. Expression changes (Log_2_FC, colour bar) of KEGG metabolic pathways active UC vs Control. (Black cell outline corresponds to pval adj <0.05; mixed linear model, Benjamini-Hochberg correction). **c,** Representative immunofluorescence of control, UC non-inflamed and. inflamed epithelium stained for EPCAM (blue), VDAC/Mitochondrial mass (green), Complex 1(red) and Complex IV (yellow). Top image scale bar 100µm, bottom image scale bar 25µm. **d,** Quantification of VDAC1 (mitochondrial mass), Complex I and IV (n=10 control and 11 UC; one-way ANOVA with Tukey’s multiple comparison test). e. UMAP visualization of clustering of epithelial cell types identified from single-cell spatial transcriptomic data (CosMx, 960 gene panel) of 222,196 cells from 157 fields-of-view (FOV) from spatial transcriptomics of 1 HC and 1 UC active patient.

**Extended Data Fig. 2.**
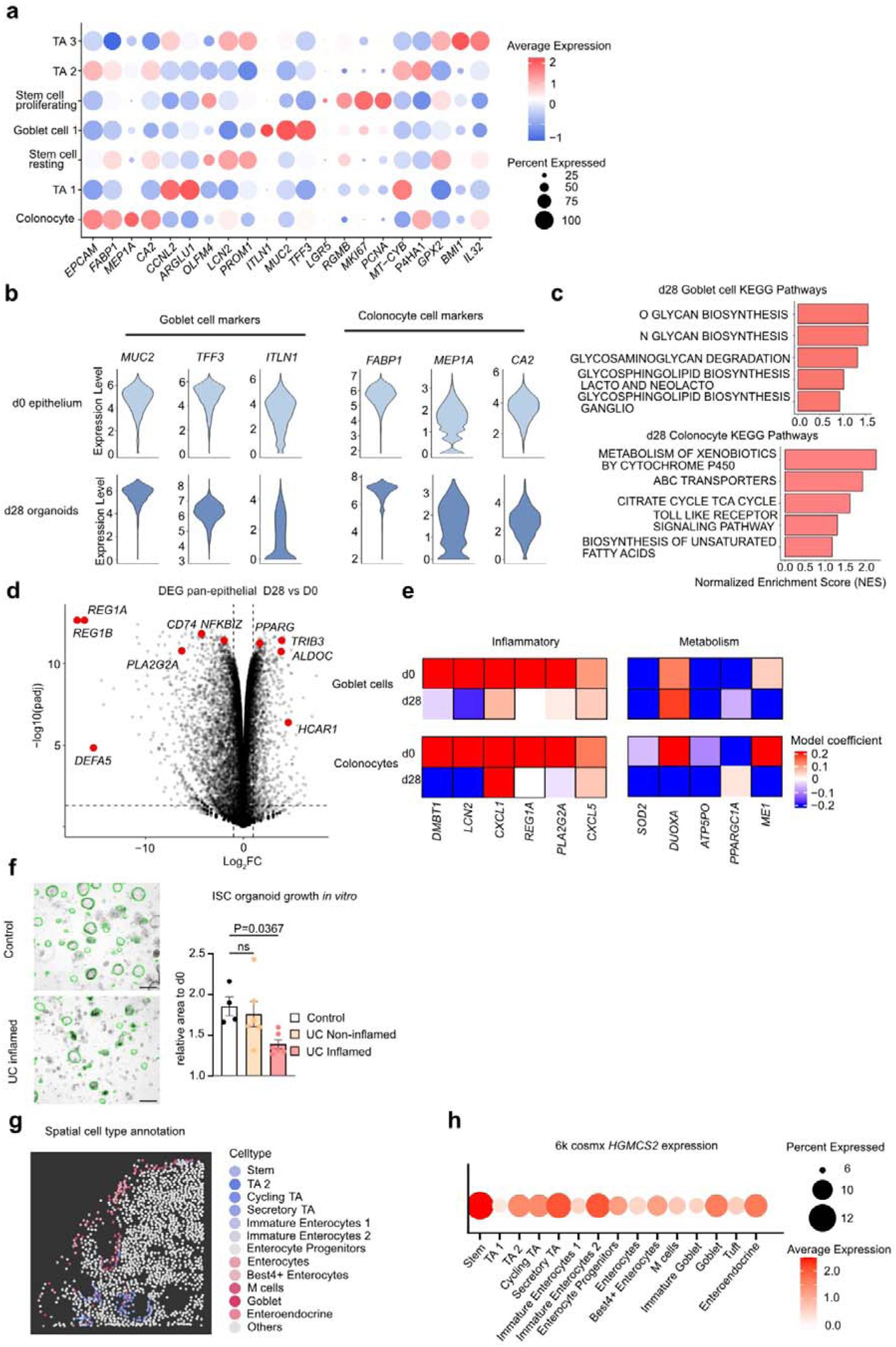
| Persistent metabolic phenotype of UC epithelial cells *in vitro*. **a,** Expression of key marker genes for each scRNAseq d28+ organoid cell type. **b,** Expression of key Goblet cell and colonocyte marker genes in goblet cell and colonocyte subpopulations derived from scRNAseq d28 organoids. **c,** KEGG pathway expression goblet cell and colonocyte subpopulations. **d,** Differential gene expression in UC active organoids vs epithelium. Pseudobulk, via edgR. **e**, Metabolic signature maintained in d28 UC organoids. Expression of key DEG inflammatory and metabolic genes in d0 Goblet cell and colonocytes and d28 Goblet cell and colonocytes. (UC active vs control, black cell outline corresponds to pval adj. <0.05, students *t* test, Benjamini-Hochberg correction). **f,** Representative images of organoid size analysis via OrganoID. Mean change organoid area over 5 days in vitro culture. (n= 4 control and 6 UC inflamed/ non-inflamed; one-way ANOVA with Dunnett’s multiple comparisons test). Scale bar 500µm. Results are presented as the mean_J±_JSEM. **g,** UMAP displaying spatial colonic cell type annotation. **h,** Spatial *HMGCS2* expression across epithelial cell types.

**Extended Data Fig. 3.**
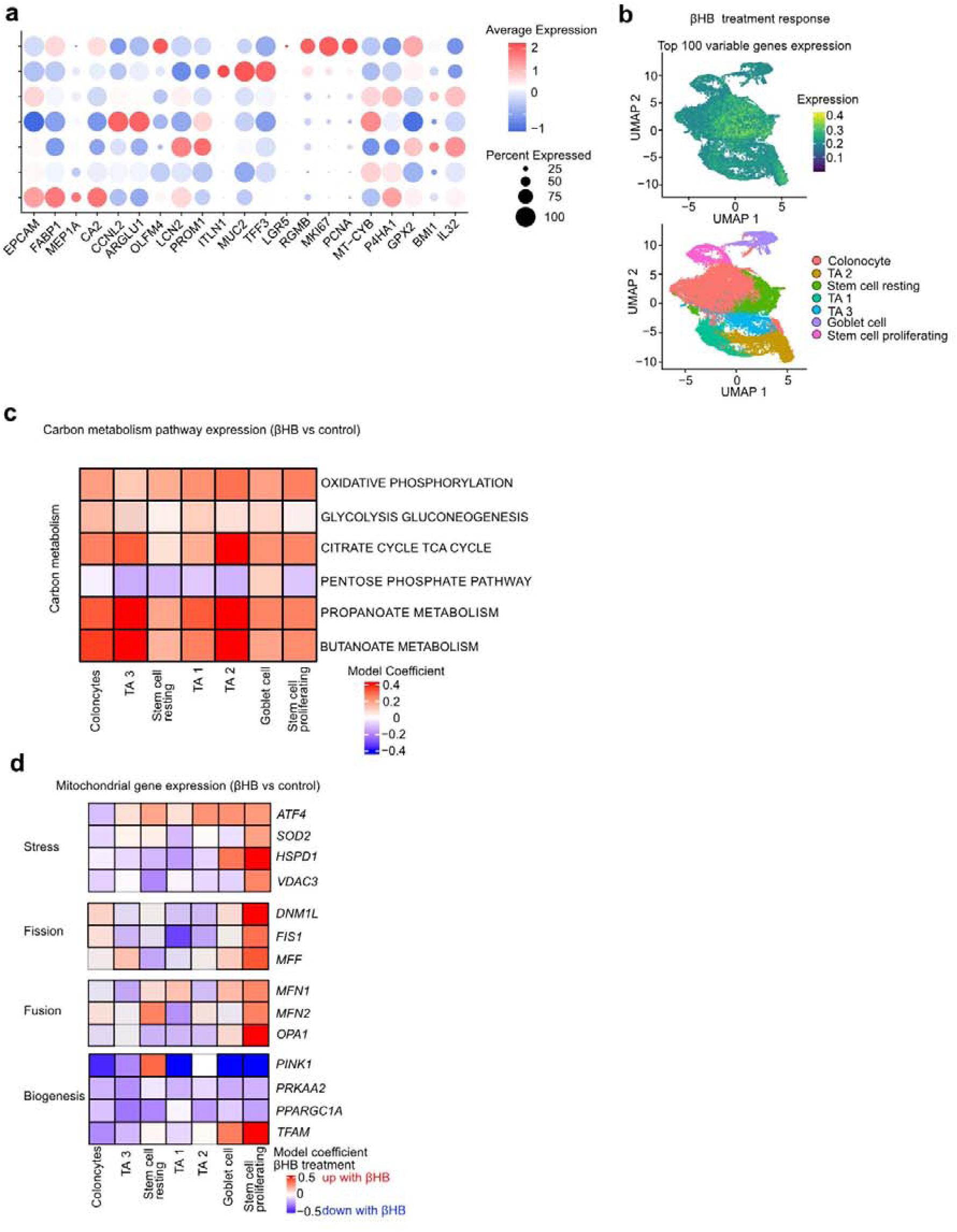
| βHB exerts transcriptomic effects across epithelial subtypes. **a,** Dot plot displaying expression of key marker genes for each organoid cell type. **b,** UMAP representation of expression of top 100 DEG via pseudobulk across βHB treated epithelial cells via scRNAseq (top). UMAP of annotated cell types bottom. **c,** Expression changes (Log_2_FC, colour bar) of carbon metabolism related KEGG pathways active UC vs Control. (Black cell outline corresponds to pval adj <0.05; students *t* test, Benjamini-Hochberg correction). **d,** Expression changes (Log_2_FC, colour bar) of key genes involved in mitochondrial stress and autophagy and biogenesis in UC organoids treated with and without βHB. (Black cell outline corresponds to pval adj <0.05; students *t* test, Benjamini-Hochberg correction).

